# “Important enough to show the world”: Using Authentic Research Opportunities and Micropublications to Build Students’ Science Identities

**DOI:** 10.1101/2023.08.17.553701

**Authors:** Lisa DaVia Rubenstein, Kelsey A. Woodruff, April M. Taylor, James B. Olesen, Philip J. Smaldino, Eric M. Rubenstein

## Abstract

Primarily undergraduate institutions (PUIs) often struggle to provide authentic research opportunities that culminate in peer-reviewed publications due to “recipe-driven” lab courses and the comprehensive body of work necessary for traditional scientific publication. However, the advent of short-form, single-figure “micropublications” has created novel opportunities for early-career scientists to make and publish authentic scientific contributions on a scale and in a timespan compatible with their training periods. The purpose of this qualitative case study is to explore the benefits accrued by eight undergraduate and master’s students who participated in authentic, small-scale research projects and disseminated their work as coauthors of peer-reviewed micropublications at a PUI. In these interviews, students reported that through the process of conducting and publishing their research, they developed specific competencies: reading scientific literature, proposing experiments, and collecting/interpreting publication-worthy data. Further, they reported this process enabled them to identify as contributing members of the greater scientific community.

## INTRODUCTION

“Every piece of data is important, but before micropublications, it was hard to get some of those smaller bits of data published…however, micropublications place emphasis on every piece of data. Every piece of data can help someone else in the scientific community.”

-Jessica, Biotechnology Certificate Student*

*All student names are pseudonyms.

Advanced academic curriculum is designed to promote talent development through the exploration of rigorous content, development of key process skills, and production of high-quality products (Olszewkei-Kubilius et al., 2018). Throughout students’ careers, their educational experiences must evolve to emphasize and develop domain-specific abilities. At higher levels of this developmental process, students need to participate in authentic tasks, guided by an expert in the field of study and designed for authentic engagement (Kettler, 2016). This principle is embedded within many advanced curriculum models, such as the Parallel Curriculum Model (Tomlinson et al., 2002). This model recognizes the value of core content and information transfer, but its unique contribution is the emphasis placed on providing students opportunities to assume the role of the practicing professional and reflect upon their role within the discipline, both now and in the future (Tomlinson et al., 2002). While the tenets of advanced curriculum are clear, implementing them in higher education institutions can be challenging, given large class sizes as well as limited time and resources. Thus, this paper chronicles the efforts of faculty at a primarily undergraduate institution (PUI) as they strive to develop talent by placing students in the role of a practicing professional and supporting the development of students’ science identities.

### Developing Science Identities

A science identity is built upon the dynamic interaction among personal characteristics (e.g., beliefs, self-efficacy) within specific social contexts (Aschbacher et al., 2010; Carlone & Johnson, 2007). As individuals develop identities, they are more likely to behave in ways that align with their identity. In turn, those behaviors strengthen their identities, forming an iterative interaction among social outcomes, personal identity beliefs, and competency development (Bucholtz et al., 2012; Gazley et al., 2014). Students with strong science identities believe themselves to be scientists belonging within the broader science community. Such students express high levels of science interest and increased intentions to pursue advanced science studies in college or graduate school (Carlone & Johnson, 2007; Merolla & Serpe, 2013; Merolla et al., 2012). Further, students with strong science identities are more likely to be employed within the science field after graduation (Stets et al., 2017). Even after controlling for alternative explanations (e.g., GPA, self-efficacy, and goal theory), researchers determined a strong science identity increases the likelihood of minority students entering the science workforce (Stets et al., 2017).

Given the positive outcomes associated with building a strong science identity, determining how to best build students’ science identity is an important pursuit. Research has examined interventions that address multiple entry points to building science identities (e.g., high school science programs; Stets et al., 2017) and identified potential constructs associated with science identities (e.g., performance, competence, and recognition; Carlone & Johnson, 2007). The current case study builds upon that work by exploring the effects of conducting authentic research coupled with coauthoring short-form, peer-reviewed micropublications on biology undergraduate and master’s students’ science identities.

Historically, providing opportunities for undergraduates and master’s students to engage in authentic research resulting in recognition from the broader scientific community has been challenging, even though this broader recognition could be key to building positive science identities. Traditional cellular biology journals require articles to make substantial advances to the field, which are often the accumulation of countless experiments, across multiple years. Thus, institutions that primarily serve undergraduates (PUIs) as well as those serving students completing terminal master’s degrees are often unable to provide research students the opportunity to see their work integrated into a publication during their time in the lab. Whether a student is still a lab member when their work is published is highly dependent upon the research project’s phase at the time they enter the lab, which is typically out of the student’s control.

While students may be authors on peer-reviewed papers, by the time these papers have been published, many of these students will have already entered their next career phase. Such publications mostly benefit the principal investigators, the institutions, and the broader scientific community, without supporting the science identity development of the students.

Recently, a new journal generated a potential solution to this problem; *microPublication Biology* is a peer-reviewed, indexed journal that provides space for smaller biological stories. The aim and scope of the journal is to “publish single, validated findings that may be novel, reproduced, negative, lack a broader scientific narrative, or perceived to lack high impact.” This journal requires articles to be peer-reviewed and adhere to standards for scientific rigor and reproducibility. While originally only publishing work on *Caenorhabditis elegans* (i.e., roundworms), in 2019, *microPublication Biology* expanded its scope to include *Saccharomyces cerevisiae* (i.e., budding yeast). Given that *Saccharomyces cerevisiae* is the key model organism used within our independent research and teaching labs, this enabled us to more frequently provide publishing opportunities to our undergraduate and master’s students at a PUI.

*microPublication Biology* provides a unique opportunity for undergraduate and master’s students (whose time in a research lab is limited) to publish their work while they are still in the lab. This opportunity could provide increased recognition of their scientific contributions, enhancing their science identities, which in turn, could positively influence their future career decisions. Therefore, the purpose of this qualitative case study is to present an assessment of (a) how we developed a system to promote undergraduate and master’s students’ opportunity to conduct and publish authentic scientific research and (b) how we leveraged that system to create a community of scientists. Collectively, this system was designed to support the development of students’ science identities by first, increasing their competencies through participation in authentic/relevant experiences and second, demonstrating how they can contribute to the broader field through micropublications.

We anchored our definition of science identity on Bernard’s (2020) conception. She synthesized myriad research articles’ definitions and approaches (Aschbacher et al., 2009; Bucholtz et al., 2012; Carlone & Johnson, 2007; Gazley et al., 2014; Vincent-Ruz & Schunn, 2018), proposing a science identity is:

> …how an individual seeks to be a scientist, which is constructed through iterative interactions with scientific social and material contexts. A person with a strong science identity would exhibit a sense of community and affiliation built by consistent extrinsic and intrinsic attitudinal factors. This sense of identity can be built by participating in relevant activities followed by categorizing oneself as a member in the scientific community.

This definition has several key components: (a) participation in relevant activities, (b) iterative interactions, (c) motivational factors (e.g., development of competence), and (d) sense of community. Each of these components are explored within this case study.

### Designing and Providing Relevant Research Experiences

To build a science identity, students must recognize they are participating in research experiences as actual scientists. However, many undergraduate science laboratory courses still rely on “recipe-driven” experiments with pre-determined, known outcomes that students strive to replicate. While these experiences support student skill development (e.g., pipetting), they do not necessarily enhance students’ deeper understanding of how actual scientific contributions are made or provide the opportunity to make such contributions (Delventhal & Steinhauer, 2020). Those lab experiences do not allow students to test original hypotheses, generate novel data, and, especially, publish their findings for the broader scientific community (Burnette & Wessler, 2003; Knutson et al., 2010).

Recently, leaders in undergraduate science education have advocated to embed authentic science learning opportunities into all students’ undergraduate lab-based courses, and many programs have heeded that call, leading to successful outcomes (e.g., D’Costa & Shepherd, 2009; Delventhal & Steinhauer, 2020; Lau & Robinson, 2009). Providing students with more authentic laboratory experiences has the potential to increase student retention in STEM, motivation, and promote science identities, better preparing and inspiring students to pursue future STEM careers (American Association for the Advancement of Science, 2011). Two main approaches can provide these experiences: (a) Course-Based Undergraduate Research Experiences (CUREs) and (b) experiences within independent Principal Investigators’ Research Labs (PIRLs). Often these experiences are considered separate opportunities; however, we conceptualized how research conducted in both CUREs and PIRLs could be integrated to create more opportunities for undergraduate and master’s students to coauthor peer-reviewed micropublications.

### Course-Based Undergraduate Research Experiences (CUREs)

First, CUREs are laboratory courses that encourage students to craft and address original research questions. Implementing CUREs in undergraduate curriculum can expand access to research experiences and more broadly expose students to authentic science (Bangara & Brownell, 2014; Esparza et al., 2020; Sun et al., 2020). Unlike a traditional protocol-driven laboratory experience, the experimental outcomes of CURE laboratories are typically unknown. Students are guided through a process that involves drafting a testable hypothesis and designing experiments to answer relevant scientific questions. While no two CUREs are the same (e.g., investigations of zebrafish development; D’Costa & Shephard, 2009; structure-function analysis of glyceraldehyde-3-phosphate dehydrogenase; Lau & Robinson, 2009), involving students in a more authentic scientific experience benefits both students and faculty. Students report a higher ability to both think and communicate like a scientist after participating in a CURE (Brownell et al., 2015). Further, students who participated in a CURE experience better outcomes in areas related to self-efficacy, self-determination, and problem-solving when compared to students who did not participate in a CURE (Olimpo et al., 2016). Faculty who teach CUREs report additional benefits, including the opportunity to connect teaching to research, positive contributions to promotion and tenure, and the opportunity to publish data collected in a CURE (Shortlidge et al., 2016). Prior to the advent of micropublications, however, data derived from a single CURE would typically constitute only a small fraction of a larger published research project, if published at all.

### Experiences in Independent, Principal Investigators’ Research Labs (PIRLs)

While CUREs provide widespread access to authentic science experiences, PIRLs provide in-depth mentorship to a smaller number of students. These experiences are high-quality and time-intensive, but they are limited in number. Like CUREs, PIRLs allow students to be immersed in an authentic research process and teach students how to think and work like scientists. The benefits of PIRLs are well-characterized; undergraduate researchers are more likely to pursue graduate degrees, and they report increased understanding, confidence, and awareness of science (Russel et al., 2007). These experiences are important in helping students clarify STEM career interest, as early exposure to undergraduate research was an important factor in retaining STEM students (Graham et al., 2013; Xu, 2015). Professional success of faculty overseeing PIRLs often depends on peer-reviewed publication of PIRL-generated data. Such publications typically comprise a substantial amount of data generated over the course of multiple years. Unless undergraduate or master’s students contributed near a project’s conclusion, by the time such work is published, it is likely these early-career scientists will have moved on to different programs or positions, thereby missing out on professional advantages conferred by authorship. Micropublications provide opportunities for students with shorter laboratory residency to see their work published on an abridged timescale.

### Combining CUREs and PIRLs

While both CUREs and PIRLs provide authentic experiences, they are often conceptualized as separate opportunities for students with different benefits and drawbacks. CUREs are more inclusive, providing all students in the course with authentic, limited experience, whereas PIRLs are afforded to fewer students, but those experiences are in great depth and often sustained across semesters. Within our case study, we built a hybrid model, connecting CUREs with PIRLs, leveraging existing resources to produce novel, important experimental results (see **Figure 1**). Data initially collected in CUREs were subsequently validated in independent labs belonging to faculty researchers with similar interests (PIRLs). This workflow has the potential to systematically involve a greater proportion of STEM students in authentic scientific discovery, leading to regular micropublications.

**Figure 1.**
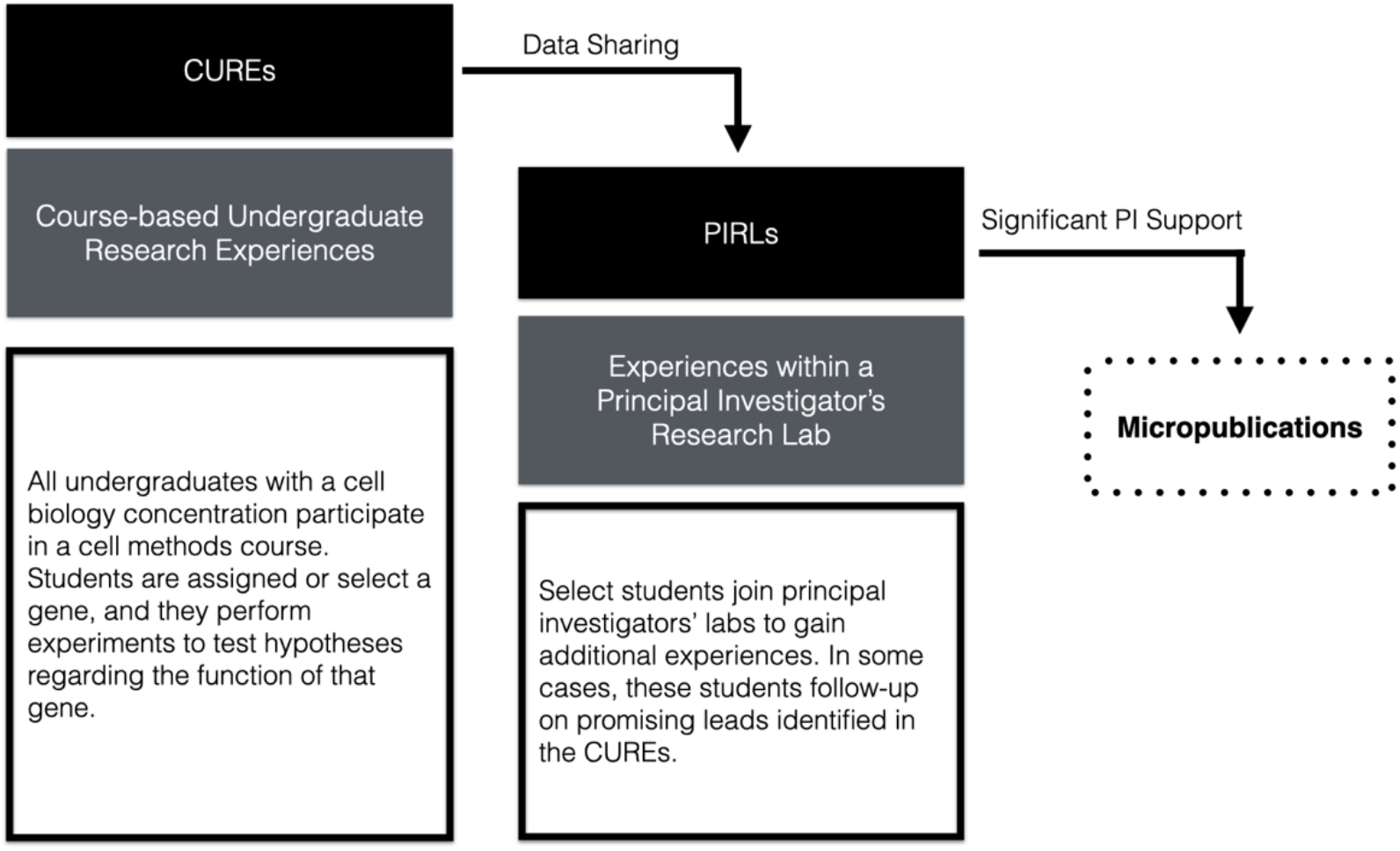
Developing a Research System: The CURE-to-PIRL Workflow.

### Building Authentic Scientific Communities

In addition to providing authentic research experiences, building a science identity requires students to develop a sense of belonging within the science community, which can be developed using multiple methods (e.g., increasing representation in featured speakers and faculty). However, there is a difference between believing you *can* be a scientist and believing you *are* a scientist. Specifically, scientists contribute to the broader community in many ways, including sharing their findings through publications and at conferences. However, those contributions – particularly publications – are often outside undergraduate and master’s students’ experiences.

When dissected, “scientific publication” is the process by which researchers disseminate experimental results to a broader academic community; the process is rigorous, typically involving multiple rounds of peer review and revisions before acceptance. To early-career science students (who have been known to complain when asked to submit a second draft of their work), this process is a complete mystery. However, providing opportunities to publish has the potential to provide multiple benefits, including enhanced student motivation and career preparedness (Cooper & Brownell, 2018; Corwin et al., 2018; Sun et al., 2020). Further, students who were immersed in research, including authoring journal papers, experienced more positive outcomes from their research (Russel et al., 2007).

Providing undergraduate students with more opportunities to publish has the potential to improve retention rates and outcomes for STEM students. Further, many graduate school admission committees and industry hiring personnel prioritize students with research experience; inclusion of a publication on a CV and being able discuss it in interviews provides an advantage to students. Publishing may allow students to see themselves not just as members of the scientific community but as active contributors as well. However, given the limited time undergraduate and master’s students spend in the lab, obtaining publications in a timely manner is challenging. With micropublications, however, students can publish their single-finding scientific stories rather than wait until a much larger, multi-figure paper is ready. These smaller publications allow students to have more ownership (autonomy) and to develop competence, providing a mutually beneficial opportunity for the lab PI, which may increase a sense of collective relatedness as they work together to accomplish a common goal.

## CASE STUDY PURPOSE

The current case study explores student authors’ perspectives as they participated in a research system, resulting in at least one micropublication. Understanding their experiences illuminates the extent to which these experiences support their science identity development. Further, understanding these students’ experiences can help professors refine the system, design longitudinal studies, and provide baseline information for other institutions seeking to implement a similar lab science program. The overarching research question of this work is: How did students’ science identities evolve through participation in authentic research experiences and publication of their findings?

## METHODS

### Participants

This study examined the perceptions of students from a public university located in the midwestern region of the United States. The university has an approximate enrollment of 13,900 undergraduates and 5,400 graduate students. Within the Department of Biology, approximately 120 bachelor’s and 18 master’s degrees are awarded annually. As of Summer 2020, 8 undergraduate (n = 2) and master’s (n = 6) students were authors of at least one micropublication. All were contacted, and all agreed to participate in a semi-structured interview (see Appendix A for protocol). Of these students, 2 identified as male and 6 as female. All were white, and, currently (as of June 2023), all remain involved in science (4 are employed as scientists in pharmaceutical or biotechnology companies, and 4 are enrolled in biomedical graduate or professional academic programs). Graduate students interviewed in this study pursued a terminal master’s degree in biology (i.e., the master’s degree was not an intermediate stage of a broader doctoral program). Students in the biology master’s program complete graduate courses and independent research in a faculty member’s research lab.

Importantly, since data collection (all data were personal interviews conducted in Summer 2020), an additional 8 students have become authors on micropublications. Of these students, 5 identified as male and 3 as female. 5 of these students identified as white, 1 identified as Asian, 1 identified as Arab, and 1 identified as Pacific Islander and Asian. In addition to the students interviewed for this report, numerous undergraduate students (*n* = 185) participated in CUREs to complete authentic pilot research studies leading to subsequent investigations to be conducted in PIRLs for inclusion in micropublications (see **Table 1**). Future work is needed to examine these additional students’ perspectives and experiences.

**Table 1.** Undergraduate and Master’s-Level Student Involvement in Micropublications.

| Micropublication Citation | Initial CUREs* | Replication and Extension in PIRLS** |
| --- | --- | --- |
| Citation to be added after blinded review. | Fall 2019 ($n = 46$) | Fall 2019 ($n = 1$) |
| Citation to be added after blinded review. | Spring 2018 ($n = 49$) | Spring 2019 ($n = 3$)<br>Fall 2019 (after reviews; $n = 2$) |
| Citation to be added after blinded review. | None. | Fall 2019 – Spring 2020 ($n = 1$) |
| Citation to be added after blinded review. | None. | Fall 2019 – Spring 2020 ($n = 1$) |
| Citation to be added after blinded review. | Spring 2020 ($n = 43$) | Spring 2020, Fall 2020 ($n = 4$) |
| Citation to be added after blinded review. | Spring 2021 ($n = 47$) | Fall 2021 – Spring 2022 ($n = 2$) |

\*Course-Based Undergraduate Research Experiences (CUREs)
\*\*Principal Investigator Research Labs (PIRLs)
\*\*\*All publications were published in *microPublication Biology*, except for one paper of similar scope, which was published in *Fine Focus*. *Fine Focus* is an indexed undergraduate-focused and managed, peer-reviewed journal.

### Research Design

To understand how students developed science identities through participating in authentic research leading to publications, we employed a qualitative case study design. A key feature of case studies is their bounded nature (Patton, 2014; Yin, 2014). Our case study is bounded by shared physical context (i.e., biology labs within a PUI), shared experiences (i.e., being a published student author), and time (i.e., published 2020 or earlier). In general, the goal of case study research is to present the inherent complexity occurring within this bounded system (Yin, 2014). Within the current study, we use this approach to determine how students evaluate and interpret their research experiences within this specific system.

### Data Collection Process

Prior to any data collection, our university’s Institution Review Board (IRB) reviewed and approved our study (Project Approval #1604061). During the summer of 2020, a graduate assistant from the Department of Educational Psychology contacted all biology student authors (*n* = 8) with information regarding participating in this case study. The student authors had no previous interactions with the graduate assistant and were not offered any incentives for participation. During the consent process, potential participants were emailed and informed that all interview data would be transcribed and anonymized before sharing with any faculty members. All student authors agreed to participate and submitted consent forms prior to the interview. Then, the graduate assistant scheduled and hosted all interviews on Zoom using the semi-structured interview protocol in Appendix A. The graduate assistant recorded and transcribed all interviews, yielding approximately 700 lines of codable data. All quotes are from personal interviews conducted during the Summer of 2020.

### Data Analysis

All transcripts were organized within Microsoft Excel files. Each conversation turn was placed in its own cell. The primary coding team was composed of a professor and graduate assistant from the Department of Educational Psychology. The coding team used Saldaña’s (2021) cyclical coding methods to code all interview data. During the first cycle of coding, they organized the data using structural codes developed from a self-determination theory lens (i.e., autonomy, competence, and relatedness). Simultaneously, the coders also recorded initial thoughts and noted participants’ high-impact phrases (i.e., in-vivo coding) within this first cycle.

As the coding progressed, they realized that the self-determination theory did not fit the data best, but rather the concept of a science identity more succinctly encompassed the data. Self-determination theory emphasizes autonomy and promotes motivation, which was present in these students’ interviews; however, students more strongly emphasized how they developed as a scientist and how they could contribute to the broader field. This caused a shift in the theoretical framework to examine how students were developing their science identities.

The coding team revisited the data to identify examples and non-examples of science competency development and belongingness within science. The science *competencies* were coded using process codes (i.e., codes that identify actions), and science *belongingness* was coded using values codes (i.e., codes that identify values, attitudes, and beliefs). Then, the coders used the transitional coding method (i.e., coding the codes), to synthesize and streamline the language across participants. During the second cycle of coding, the coders used focused coding to ensure the codes could be applied throughout the data set without needing further adaptations.

Several methods were used to ensure reliability and validity within the qualitative data analysis (Merriam & Tisdell, 2015). Specifically, the coding team consisted of multiple coders, which promoted reliability, and they piloted different coding methods to promote validity. The team kept an audit trail of their coding decisions, and within this manuscript, all findings are supported by participants’ quotes, using rich and thick descriptions.

## RESULTS

As this case study evolved, we began to see these authentic research experiences culminating in micropublications as more than a mechanism to increase student motivation, but rather as an approach to support students’ science identity development. The overarching findings align with Bernard’s (2020) definition and demonstrate the importance of (a) engaging in relevant activities and (b) joining as well as contributing to the broader scientific community.

**Finding #1: Engaging in relevant activities (i.e., understanding scientific literature, proposing research questions/experiments, collecting publication-quality data, and interpreting data) were important learning experiences for developing students’ science identities**.

Building a science identity requires students to participate in authentic science; however, authentic science experiences are not often presented within undergraduate education. Archer et al. (2010) describes this as the misalignment between school science and real science. Students know when they are simply replicating old (“recipe-driven”) experiments with known, correct answers. However, engaging in authentic science is challenging given the vastness of the field; it is first necessary to acquire new language, understand complex content, and develop experimental techniques before engaging in authentic science. Within this research-to-micropublication workflow, students were scaffolded into authentic science experiences in a smaller, contained project, positioning students to develop authentic language and research skills, albeit in a narrow sphere.

### Understanding Scientific Literature

To make meaningful scientific contributions, students needed to understand the state of the field. While textbooks provide a conceptual summary, reading contemporary research is necessary to understand both current discoveries and experimental methods leading to specific conclusions. Several students discussed the difference between reading science textbooks and reading scientific research articles, noting scientific research articles included specialized vocabulary and notations that were initially foreign to them. Beyond language complexity in primary literature, students also described challenges with understanding why scientists did a certain experiment, how that experiment yielded the data, and how the data was used to derive specific conclusions. Jessica summarized how she initially experienced these challenges, but, how, through engaging in microprojects and contributing to micropublications, she developed these competencies:

> I definitely understood my course content. I did very well in my classes. But that’s very different than going into a research lab and understanding what they’re trying to study. When I first started in the lab, I felt like I knew nothing because even though I had taken two and a half years of college classes, being able to read a research article on what we were studying was nearly impossible. They use different jargon. Then, looking at their data and trying to figure out how they got those data from the experiment they did, I couldn’t understand what experiments they were performing. I felt like I knew nothing when I started. It was very much a slow process to start to understand how to read an article and how to understand what they did, why they did it, and understanding the foundation of our research. But now I feel very, very confident reading articles. I can look at the data critically, and I can understand what they did and understand the controls that they used. But that definitely was not something that I felt confident in at first…If you can write an article and if you can proofread one, you can totally read and understand one.

Brianne echoed this perspective, “I’ve become more comfortable working through papers and trying to figure out what they were doing. Expressing my own science and also understanding other peoples’ science has improved.” Further, Elizabeth suggested micropublications could serve as a gateway for new students to understand scientific publications in a manageable size, saying “this type of format seems so helpful and usable…a steppingstone for people to read this type of work and get things from it.”

### Proposing Research Questions and Experiments

Being able to read and understand existing literature positioned students to be able to propose new experiments. One of the eight students independently developed the primary initial research question leading to a micropublication. Several other students proposed follow-up experiments for projects leading to micropublications that addressed research questions.

First, Josh proposed the primary research question leading to a micropublication after he connected knowledge from external publications with another lab student’s project. Josh brought this connection to the PI, who encouraged him to test it. Josh reflected upon the experience, “That led me down my own path of doing my own experiments to see if this pathway has any correlation with what we’re working on. It turns out it didn’t, but it was still when I began questioning things.” While most students were not responsible for the primary question, many students assumed more responsibility for proposing the next experiment. For example, Jessica described:

> Usually, we’d get a piece of data, and we’d consider 10 experiments that we needed to do. And the PI would ask: “where do you think we should go?” I would say, “I think the first experiment we need to do is this one, because that will tell us this, and then, we wouldn’t even need to do this one.”

### Collecting Publication-Quality Data

For micropublication-scale projects, collecting publication-quality data to address research questions and follow-up experiments is an important objective from the project’s genesis. Generally, some data collection competencies can be developed without the end goal of a publication, and within any lab experience, a certain amount of troubleshooting is necessary. Grace provided an example of how she needed to troubleshoot to ensure a successful experiment:

> I was doing this experiment where I had to grow some yeast in liquid media. I was measuring their growth rate over time in a certain drug. I noticed the drug was automatically killing them [when added in a later step], so I suggested an adjustment: growing them in the drug first, before starting the experiment.

Working towards publishable data, however, required an additional layer of understanding and influenced how students approached implementing and reading protocols to ensure replicability. Amanda discussed her struggle to replicate the data, yet “…it was a big confidence boost to be able to sit down and do it over and over and over with zero issues because I had been optimizing this protocol for three months…” When publications are not the key outcomes, there is not the same level of motivation to “make every gel a figure.”

> Jessica specifically described the relationship between protocol specificity and replicability needed for publications.
>
> In [previous] classroom labs, I learned a little bit about how to pipette and how to read a protocol and follow steps, just like a recipe. But being able to be so consistent that if I ran an experiment this week, I could replicate it again next week and get the same result, that definitely took work to get there. It was a lot of training and understanding the techniques. Is this a step that has to be done in a very specific way in order for it to work? What needs to be…considered when you’re looking at a protocol? Every step of the protocol is important, but there are some [ambiguous] steps. [For example,] “add four things.” Do you have to add them in a specific order for it to work? So definitely over time, that ability improves significantly to understand how to do a protocol in a really effective way…a way that is able to be replicated and give you really strong data.

If students were simply developing competencies in specific techniques and use of equipment (e.g. in traditional science laboratory class), they may not have developed this deep recognition of the importance of being able to replicate results. The system of data collection promoted this understanding by involving initial undergraduate lab studies and then replicating those findings in specific PI labs.

### Interpreting Data

After optimizing protocols and replicating results, students needed to be able to translate data into new understanding about cellular processes, and that translation required practice. Elizabeth described the “aha moment” where she was able to understand the “cellular mechanisms behind the physical data.” Similarly, Brianne described in more detail a specific instance when she realized she was able to use data to understand a cellular process:

> One moment is when we saw an unusual result in my experiments. And instead of [the PI] telling me what he thought was happening, he asked me what I thought was happening. I told him, and it was exactly what he was thinking! Before that, I would have never been able to think about the cellular level and what was going on and why we were getting this result. But in that moment, I was able to work through my experiment, the problems we were having, and actually connect the two, to connect the results to what was going on in the cell.

Developing these authentic competencies were essential in establishing students’ self-efficacy for doing science, as Grace suggested:

> …being in an actual lab, you have more of an opportunity to do things on your own, and you kind of get things done faster too, because you’re not relying on a group or like a professor to teach you something…It’s easier to learn things on your own and see data for yourself. And you kind of feel more accomplished because it’s the data that you produced on your own instead of like, your group is progressing through like a recipe protocol and seeing the data at the end.

**Finding #2: Joining the scientific community was important across levels of dissemination, including within the lab, at conferences, and within micropublications.**

In addition to engaging in authentic science, building a science identity requires seeing oneself as a member of the in-group. Students within these labs identified multiple experiences that helped them feel like they belonged, both at the level of the individual research lab and within the broader scientific community through their micropublications.

### Finding One’s Voice in the Lab

Many students initially felt hesitant to ask questions or suggest potential solutions because they were uncertain of how the group would perceive them. However, being able to use one’s voice is a key aspect of being a part of the scientific community, and within these labs, students were able to find their scientific voice over time due to deliberate encouragement from the PIs. Brianne described how she was able to find her voice in one-on-one mentorship meetings with the PI:

> We would have weekly meetings with [the PI] in which I would talk about my experiments and what was going wrong and what was going well. In those meetings, it was easy to talk about problems we were seeing. [The PI] would suggest things and then ask me if I had any ideas, and sometimes I didn’t, but sometimes I did. It really helped that [the PI] was willing to hear what I was saying, and it made me feel comfortable speaking up. At the beginning of the lab, I probably wasn’t comfortable speaking up and suggesting things, but as I got more comfortable, I was able to look at my experiments and think of new ideas for it. And it was definitely a space in which we were encouraged to come up with ideas and suggest things.

Beyond providing ideas for troubleshooting and experiment planning, students also felt comfortable revealing moments in which they were uncertain, as Josh described:

> I feel like I can express my ideas really freely…it’s a bit of the culture that [the PI] has, where questions are seen as the curiosity, that they are not something to be embarrassed about…[the PI] makes it okay to ask questions, no matter what they are.

### Presenting to the Broader Field

In addition to contributing to the local lab community through questions and ideas, these students also recognized and valued the opportunity to contribute to the broader field. Specifically, they practiced communicating their findings in both formal and informal settings, including lab meetings, conferences, theses, and, eventually, micropublications. Collectively, these experiences helped them develop fundamental scientific competencies and enhanced their confidence in their own abilities to contribute to the broader field. When interviewed, they emphasized their growth in scientific communication and recognized the importance of communicating complex results in a way that others could understand. For example, when Grace was asked how she has grown as a scientist, she described how she became more comfortable presenting:

> …So [previously] I gave a one-hour presentation, and I got so nervous. I could not speak at all. Now, while presenting my thesis, I spoke without hesitating. I knew exactly what was going on, and I just felt so much more confident in myself, in what I was saying and how I was expressing my data.

Similarly, Shannon described her growth:

> When I first started in the lab, I felt less confident speaking up…but working on my thesis, I had to be able to write and talk about things in a way that would make sense to someone who isn’t a biologist, so knowing that I both understood what I was talking about and was able to explain it to someone who might have no idea, in a way that they could also understand, really showed me that I grew.

### Publishing in a Scientific Journal

In addition to recognizing their own growth in communication skills in the local university setting, they also grew to appreciate overarching science communication principles, including the variety of methods and formats employed in their field and the importance of conciseness, all of which help readers better understand their work. Participating in both the initial development process and the subsequent revise-and-resubmit process enhanced students’ ability to communicate and enabled them to gain an improved understanding of the publishing process. Grace reflected:

> …it’s made me realize how hard the writing can be. It’s not like you write it and that’s what you publish. You have to write it and edit it, like six times, before it actually can get published and approved…There’s just a lot more that goes into publishing than I expected. The people that review your manuscript are very picky. They know the field well, so you have to make sure you know what you’re talking about too.

While some of the manuscripts required only minor revisions, some required considerable adaptations. Specifically, Rob discussed going back and forth with reviewers on the need to include a statistical analysis: “they weren’t happy with the statistics…And so we had to redo it and figure stuff out…by the time we finally got it in, and it was published, we were like, okay, thank God….” Rob followed up with a recommendation for the publishing process:

> …there should be a standardization of stuff that [reviewers] should not ask and stuff that doesn’t need to be done… like rules to cover when basic statistics aren’t necessary or what types of labels are necessary, or other little details. That should be just general across the board. I think that would make a lot of people’s lives easier.

While not all the micropublications required such significant revisions, the other students were aware of the challenges associated with this project, even if their own projects were being published with greater ease. Thus, being exposed to everyone’s experiences helped all students see the nuance in the publishing process. Elizabeth described that the publication process taught her:

> …that’s not the only way to do it…I always thought of [scientific writing] as one cookie-cutter mold. And the truth is that there’s a lot of different ways. Even individual journals may request different sections that I never knew existed… [I also learned] how to properly word things in a way that is most helpful to the reader.

As students reflected upon the publishing experience, they were able to identify specific communication skills they built, but perhaps even more important was the way they discussed the impact of the publication, as Brianne stated:

> I have never been confident in myself and what I do, so to see my work published by a scientific journal was really, really cool. And it gave me this confidence boost, like “you do know what you’re talking about sometimes.” That was definitely reassuring.

Similarly, Jessica described:

> It’s really very validating to know that a reviewer or a journal thought that your data and your research was *important enough to show to the world*. And that they thought you wrote well enough and concisely enough and clearly enough to convey the data in a way the general scientific community would understand.

## DISCUSSION

This study explored the impacts of engaging undergraduate and master’s student scientists in small-scale research projects leading to micropublications. Participant reflections indicate these activities enabled students to assume the role of practicing professional and supported the development of their science identities, providing a domain-specific example of advanced academic curriculum development at the college level. Further, the CURE-to-PIRL workflow addressed a common issue within PUIs, specifically that there are typically not enough spaces in authentic research labs for all students to have opportunities to meaningfully engage in the research process. Collectively, this project provides one replicable example by which advanced curriculum may be implemented in institutions of higher education, including PUIs.

While students discussed specific moments throughout their lab experiences, their experiences must be considered as a sustained interaction with the scientific community. Most of these students participated in CUREs, followed up on data from other CUREs, and joined PIRLs for at least two years. Building a science identity is not likely to be the result of a single moment; rather, these identities are developed through continuous engagement with the iterative nature of science. To demonstrate this iterative relationship, we developed the Applied Model of Science Identity. We decided to title the model “applied” because, within all interviews, students discussed the *applied* nature of their scientific identity, including specific competencies (e.g., reading scientific literature) and specific contributions to the community (e.g., disseminating findings in micropublications). The Applied Model of Science Identity (See **Figure 2**) depicts this iterative, cyclical nature: as students develop their competencies, they are then able to contribute to the community, which enhances their identity, leading to the desire to learn more and develop additional competencies.

**Figure 2.**
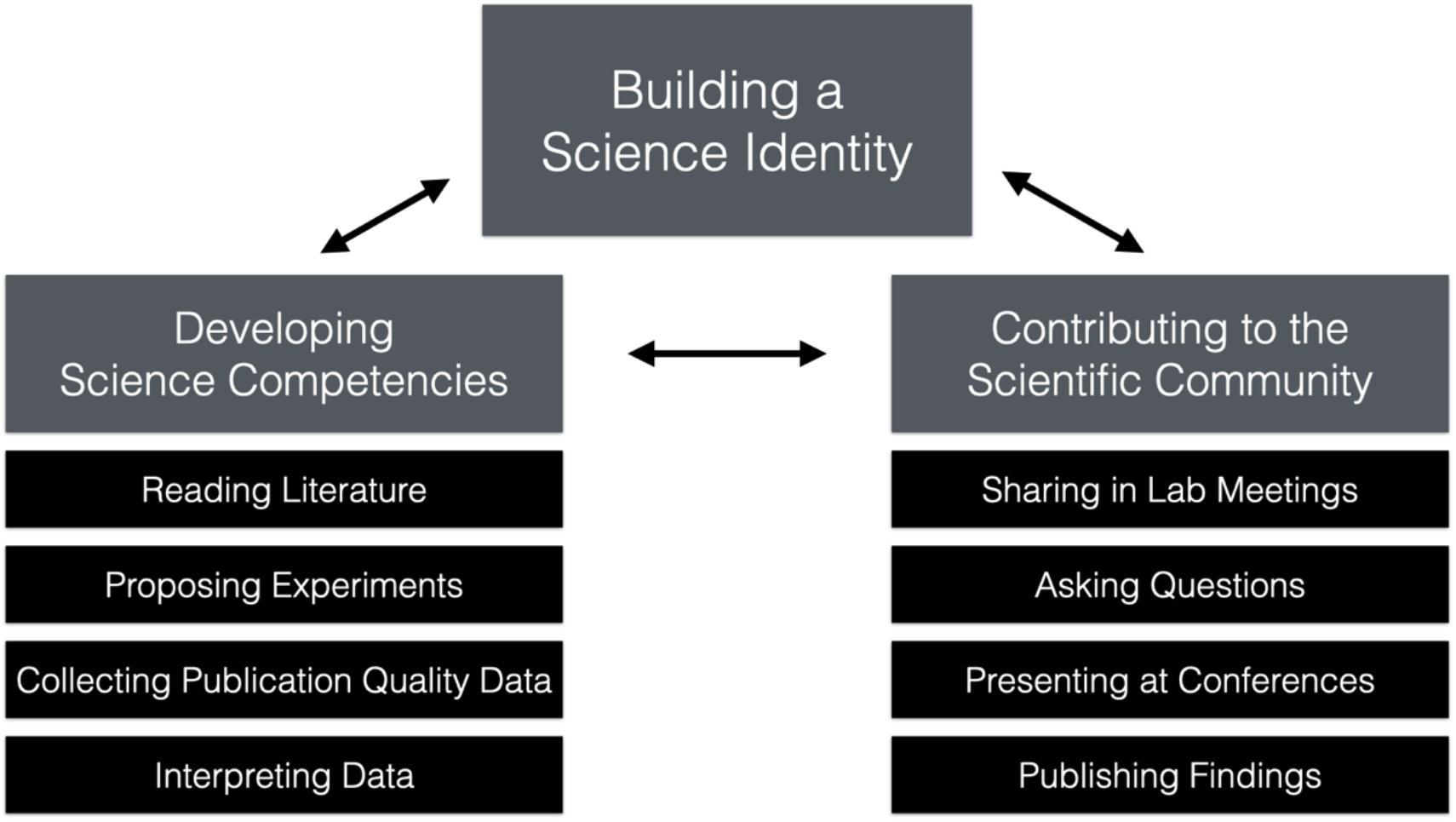
The Applied Model of Science Identity.

The Applied Science Identity Model is anchored on previous work. Specifically, Carlone and Johnson (2007) proposed a science identity model for Black women scientists through a series of interviews. The Carlone and Jackson (2007) model included competence, performance, and recognition; the recognition component was dissected into three types of science identities: altruistic, research, or disrupted. Further, they conceptualized the effect of other identities (e.g., race, gender) on participants’ science identities.

Our model represents a more generic, early science identity framework. We combined Carlone and Johnson’s (2007) “recognition” and “performance” components into “contributing to the scientific community,” which loses some of the richness of their model. Conversely, we emphasized and analyzed the competencies the undergraduate and master’s students identified. In this way, the Applied Model of Science Identity delineates specific objectives and opportunities that science programs should prioritize to build students’ science identities.

Taken together, our results demonstrate that engagement in research projects that culminate in peer-reviewed micropublications facilitates scientific identity development for undergraduate- and master’s-level trainees. Our work strongly suggests that participating in a large-scale, multi-year project is not necessary to realize desirable psychosocial benefits for early-career scientists. Further, because projects leading to micropublication are by their nature smaller in scope than traditional research projects, faculty mentors with limited time or resources may be empowered to provide authentic research experiences culminating in peer-reviewed micropublication for trainees. Such micropublications may reasonably be completed during early-career trainees’ limited research lab residency, providing positive personal and professional outcomes while they are most beneficial. Moreover, faculty at a range of institution types and on a spectrum of funding levels may find the CURE-to-PIRL workflow for piloting, validating, and publishing meaningful scientific results a useful model for meeting or enhancing their teaching, mentoring, and research objectives.

### Limitations and Future Research

While this case study demonstrated how combining authentic research with publication opportunities can enhance students’ competencies and promote belonginess, it represents just the beginning of a more robust research agenda. The limitations of the current study provide inspiration for future studies.

### Development of Narrow Technical Knowledge

Interviewed students recognized that most of their growth was in a specific area of biology, using specific techniques on a specific model organism. For example, Grace described, “my knowledge is less broad. Coming from a yeast lab, I was focusing on one particular area of cellular processes. My knowledge is deeper, but it’s more focused on a specific area instead of broad topics.” Several students also wanted more information and experience working with other organisms and techniques, as Brianne stated:

> The main thing I didn’t learn in the lab that could help me is mammalian cell culture. We just worked with yeast and bacteria, which I learned a lot about, but I definitely want to learn more about mammalian and human cell culture for my career…it would be good experience to have.

These students recognized that their experiences prepared them to be scientists, but scientists in a specific space with a specific specialty. Future work may be performed to consider how to optimize the breadth-to-depth ratio during early-career scientific training.

### Integration of CURE Students and Students Working in Additional PIRLs

All interviewed student authors were members of PIRLs and worked with a limited number of faculty. Nearly 200 students in CUREs, however, contributed initial pilot data leading to micropublication projects. These CURE projects were authentic research projects that generated novel findings that were refined and replicated in a PIRL. Thus, CURE students did not always see the final paper. Future work should conceptualize how to integrate these students into the publication process or at least make them more aware of their role in the workflow. As disseminating research as micropublications becomes more common, it will be important to evaluate the experiences of student coauthors in a range of research labs and at multiple institutions. After completing and analyzing interviews for this study, students in at least two other research labs at our institution have contributed to micropublication submissions. Indeed, one objective of the current study is to share the benefits of and provide a roadmap for other investigators to disseminate early-career trainee work as micropublications.

### Consideration of Multiple Identities

One final limitation is the singular focus on science identity, rather than considering how all identities interact, as Carlone and Johnson’s (2007) work discussed. In the future, it will be important to purposefully recruit and longitudinally research students with diverse identities to determine how their other identities influence and interact with science identities. Additional systems could work to integrate cultural capital and identities into students’ science identities. Despite these limitations and future opportunities, this current case study illustrates the potential of a CURE to PIRL workflow to produce high-quality, rigorous micropublications, demonstrating to students that their work is “important enough to show the world.”

## Appendix A. Semi-Structured Student Interview Protocol

### Section 1: General Lab Experiences

- First, tell me a little about what made you want to pursue a science degree.
- Tell me a little about your course lab experiences.
- How did you find the [the PI’s] Lab?
- What made you want to join?
- Tell me a little about your initial impressions working in the [the PI’s] Lab.
- How did your impressions evolve over your time in the [the PI’s] Lab?
- What was the most memorable moment in the [the PI’s] Lab?
- How was your time in the [the PI’s] Lab different from your course lab experiences?

### Section 2: Developing Autonomy

- Do you ever make suggestions regarding your experiments, either adaptations to protocols or how to troubleshoot problems? If yes, can you provide an example?
- To what extent did you contribute to your publication? How much did you write?
- To what extent do you feel free to express your ideas in the lab or lab meetings?

### Section 3: Developing Competence

- How have your skills as a scientist changed throughout your time in the lab?
- Can you provide a specific example of an area in which you grew?
- How has your content knowledge changed?
- What grade would you give your scientific writing? Has it changed across the time you have been in the lab?
- What grade would you give your scientific thinking? Has it changed across the time you have been in the lab?
- What was it like to work on a micropublication?
- How did you feel when you found out it was published?
- Is there anything you wish you had learned? Or something you still want to develop?

### Section 4: Developing Relatedness

- How connected to you feel to the other members of the lab?
- How would you describe your relationship with [the PI]?
- How connected do you feel with other scientists outside the lab?
- Do you fit in with the broader scientific community? Why do you think that?
- Have you attended any conferences? How would you describe your experiences at these conferences?

### Section 5: Future

- What would you change about your lab experiences?
- Where do you think you will go from here?
- What are your career aspirations?
- Did your lab experiences influence your thoughts about the future? Can you explain?
- As a non-biologist, I may have missed some important aspects of your biology experiences. What questions should I have asked that would have captured these important experiences?

